# Role of Dynamical Instability in QT Interval Variability and Early Afterdepolarization Propensity

**DOI:** 10.1101/2025.06.11.659221

**Authors:** Daisuke Sato, Bence Hegyi, Crystal M Ripplinger, Donald M Bers

## Abstract

Beat-to-beat variability of the QT interval (QTV) is a well-established marker of cardiac health, with increased QTV (> 5 ms) often associated with a higher risk of arrhythmias. However, the underlying mechanisms contributing to this phenomenon remain poorly understood. Recently, we showed that cardiac instability is a major cause of QTV. Early afterdepolarizations (EADs) are abnormal electrical oscillations that occur during the plateau phase of the cardiac action potential (AP), often arising when the membrane potential becomes unstable. In this study, we use a physiologically detailed computational model of rabbit ventricular myocytes with stochastic ion channel gating to investigate the relationship between QTV and EAD propensity. We found that increased AP duration (APD) variability, which serves as a surrogate for QTV on the ECG at the single-cell level, can arise even in the absence of apparent EADs, driven by intrinsic dynamical instability. As the cellular state approaches the threshold for EAD generation, small perturbations in membrane voltage are amplified, leading to increased APD variability. The phase-plane analysis in the voltage-calcium channel inactivation space demonstrates that proximity to the EAD-generating basin of attraction strongly influences repolarization variability, establishing a mechanistic link between QTV and EAD propensity. Furthermore, we observed that QTV increases at longer pacing cycle lengths (PCLs), distinguishing it from alternans-associated APD variability, which increases at shorter PCLs. These findings suggest that increased QTV may serve as an early indicator of arrhythmic risk before the manifestation of EADs, potentially offering a critical window for preventive intervention. Our results provide novel insights into the fundamental mechanisms underlying QTV and its potential role in arrhythmia prediction.

**Significance statement:** This study investigates the relationship between beat-to-beat variability of the QT interval (QTV) and the propensity for early afterdepolarizations (EADs), abnormal electrical oscillations linked to life-threatening arrhythmias. Using a computational model, we show that increased QTV can precede apparent EADs, driven by inherent dynamical instability of cardiac cells. As cells approach an arrhythmia-prone state, small membrane voltage fluctuations are amplified, increasing repolarization variability and thus QTV. Furthermore, QTV increases with slower heart rates, distinguishing it from alternans, another type of instability arising at faster rates. Thus, increased QTV may serve as an early warning signal for arrhythmia risk, potentially enabling preventative interventions. Our results provide novel insights into the fundamental mechanisms underlying QTV and its potential role in arrhythmia prediction.

## Introduction

Early afterdepolarizations (EADs) are abnormal electrical oscillations that occur during the plateau phase of the cardiac action potential (AP) (1-13). EADs arise from complex interactions of ion channels and membrane potential in cardiomyocytes, often under pathological conditions. These EADs are typically associated with a prolongation of the AP duration (APD), which can be caused by a reduction in repolarizing potassium currents (e.g. I_Kr_, I_Ks_) or an augmentation of depolarizing currents such as the L-type calcium (Ca^2+^) current (I_CaL_), late-sodium current (I_NaL_), or inward sodium-calcium exchanger (NCX) current. The prolongation of the AP promotes the reactivation of the L-type calcium channels (LTCC), resulting in the oscillatory behavior of the AP. In this study, we focus primarily on phase-2 EADs due to reactivation of LTCC, which manifest as oscillations during the plateau phase of the AP. For the purposes of our analysis, an EAD is quantitatively defined as a positive deflection in membrane potential (dV/dt > 0) after the initial upstroke and before complete repolarization to the resting membrane potential.

The membrane potential is inherently fluctuating that arises from various sources, including ion channel gating, cellular metabolism, and even thermal noise. The AP plateau voltage is a delicate balance of multiple small inward and outward currents, and thus is vulnerable to relatively modest variations in any of those, thereby influencing APD. In our previous study, we demonstrated that stochastic fluctuations can modulate EAD dynamics (14). Stochastic fluctuations can either promote or suppress EADs especially near the bifurcation points.

The beat-to-beat variability of the QT interval (QTV) on electrocardiograms (ECGs) is a well-recognized marker associated with cardiac health (15-23). In general, QTV tends to increase with deteriorating cardiac health and is observed in various cardiac pathologies, including heart failure (21,24,25) and long QT syndrome (16,26,27). However, the underlying mechanisms contributing to this phenomenon remain poorly understood. Specifically, it is unclear how changes at the cellular level, such as altered ion channel function, manifest in increased QTV at the whole-heart ECG level. There are many thousands of ion channels in a cardiomyocyte. According to the law of large numbers, fluctuations at the whole cell current level are limited and thus result in small variability at the cellular level action potential. In addition, cells are electrotonically coupled via gap junctions in tissue (28), further reducing the variability. Thus, stochasticity alone cannot fully explain the large QTV observed in ECGs. How, then, can such substantial variability arise at the ECG level? Our previous study has shown that dynamical instability can amplify the small fluctuations occurring at the channel level (29). Dynamical instability can also lead to EADs (7-9). In this study, we will show that QTV increases even in the absence of apparent EADs and establish a mechanistic link between QTV and propensity for EADs.

## Materials and Methods

We used a physiologically detailed rabbit ventricular myocyte AP model from our previous study (14) which is based on the Mahajan *et al*. model (30). Modifications from the Mahajan *et al*. model include increasing I_CaL_, decreasing K currents (I_Kr_, I_Ks_) to reduce repolarization reserve, and increasing the window current by changing gating kinetics to promote reactivation of LTCCs. In the absence of stochastic noise, the model exhibits normal APs, periodic EADs, alternating EADs, and even chaotic EADs (7).

These behaviors can be modified by introducing stochastic noise (14); stochastic fluctuations can either promote or suppress EADs (Fig. 1A-B), as some EADs are highly sensitive to small changes in membrane potential.

**Figure 1.**
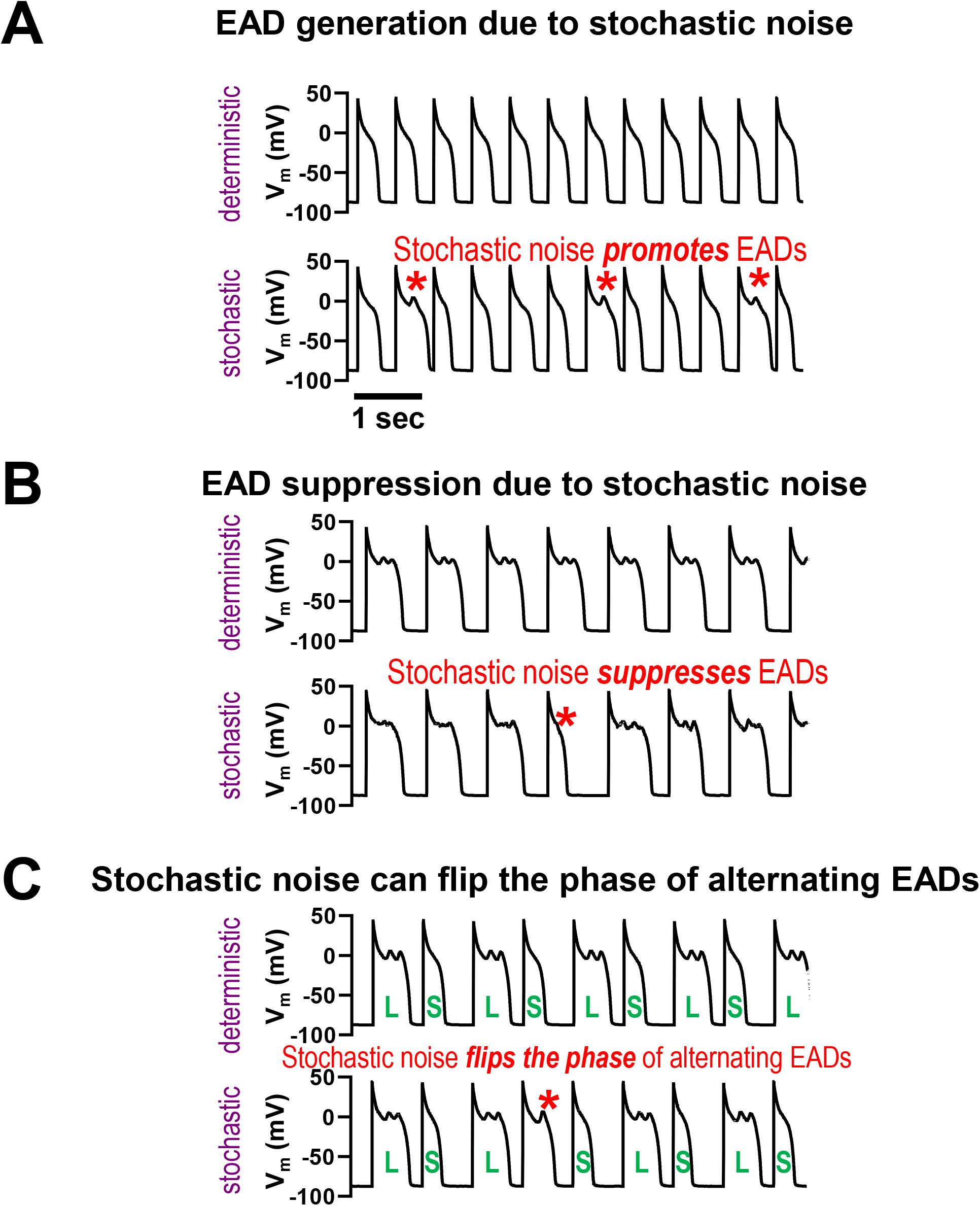
Stochastic fluctuations can promote or suppress EADs. (A) Example of EAD generation by stochastic fluctuations (B) Example of EAD suppression by stochastic fluctuations (C) Stochastic fluctuations can also flip the phase of alternating EADs, where normal APs and EADs occur alternately.

To introduce stochastic noise into the action potential, stochastic gating was implemented for the gating variables of I_Na_, I_CaL_, I_Kr_, I_Ks_, I_to,f_, and I_to,s_. LTCC is described by a Markovian model, we simulated it directly without further approximation. For the I_Na_, I_Kr_, I_Ks_, I_to,f_, and I_to,s_ channels, stochastic gating was implemented by incorporating noise terms into the gating variable equations, as described by the Langevin equation as follows (31,32). Stochasticity was applied to the gating variable (denoted as x) of the channels. The deterministic version of the gating equation is given by:

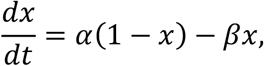

where *x* is the gating variable, *α* is the opening rate and *β* is the closing rate. The stochastic version of this equation is:

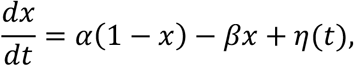

where *η*(*t*) is the noise term, which satisfies the following correlation function:

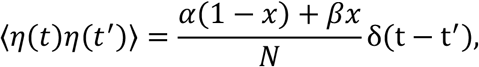

where *N* is the number of channels. Thus, the magnitude of the fluctuations decreases with an increasing number of channels, proportional to 1/√*N*. For example, if *N* is reduced or increased tenfold, the fluctuation magnitude changes by a factor of approximately 3.16. Importantly, the qualitative behavior of the results remains unchanged across a wide range of *N*. In this study, we set *N* = 100,000, which is within a physiologically relevant range (33-37). In addition, simulations with N=10,000 and *N*=1,000,000 channels were performed to assess the impact of noise strength.

The ordinary differential equations (ODEs) describing the AP model were solved numerically using the forward Euler method with an adaptive time step, ranging from 0.01 to 0.1 ms. The model was paced for at least 1,000 beats to ensure steady-state conditions.

The QT interval, measured from the start of the Q wave to the end of the T wave on the ECG, reflects the duration of ventricular repolarization. Beat-to-beat QTV, therefore, represents the fluctuations in ventricular repolarization time over successive heartbeats. In this study, the APD, which approximately corresponds to the QT interval at the cellular level, and its variability were analyzed. APD variability was quantified as the standard deviation of APDs measured over 10,000 steady-state beats.

To investigate whether the single-cell findings translate to multicellular tissue, two-dimenational (2D) tissue simulations were performed. The tissue was modeled as a 100 × 100 grid (1.5 cm × 1.5 cm) of coupled cells, using the same AP model used as in the single-cell simulations. The AP in tissue was described by the reaction-diffusion equation:

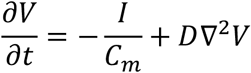

where V is the membrane potential, C_m_=1 μF/cm^2^ is the membrane capacitance, and *D* is effective diffusion coefficient, set to 0.001 cm^2^/ms. Pacing was applied at one corner of the tissue. Pseudo-ECGs were computed by summing transmembrane currents over a region of the tissue. The QT interval was measured from the pseudo-ECG. Beat-to-beat variability of this QT interval was quantified as its standard deviation over 500 steady-state beats.

## Results and Discussion

### APD variability increases even without apparent EADs

EADs are known to exhibit complex dynamics under certain conditions (7-9), typically at intermediate pacing cycle lengths (PCLs), leading to significant APD variability even in deterministic models (Figs. 2A and 2B, intermediate PCLs). In contrast, periodic APs or periodic EADs exhibit no APD variability (Figs. 2A and 2B, short and long PCLs, respectively) in the absence of stochastic noise. However, ion channels are inherently stochastic, and fluctuations in the membrane potential can significantly alter the behavior of EADs. Stochastic fluctuations can either promote (Fig. 1A) or suppress EADs (Fig. 1B) as we reported before (14). Stochastic fluctuations can also flip the phase of alternating EADs (Fig 1C). These effects happen especially when EADs are sensitive to small changes in membrane potential, as the stochastic noise can effectively push the system across these bifurcations, resulting in transitions between EAD and non-EAD states, or modifications of EAD morphology and timing. These irregular EADs and APs contribute to increased APD variability (Figs. 2C and 2D). In Fig. S1, we showed regions where EADs arise primarily due to deterministic chaos (Fig. S1A), where they are due to stochastic noise (Fig. S1B), and where EADs are absent (Fig. S1C).

**Figure 2.**
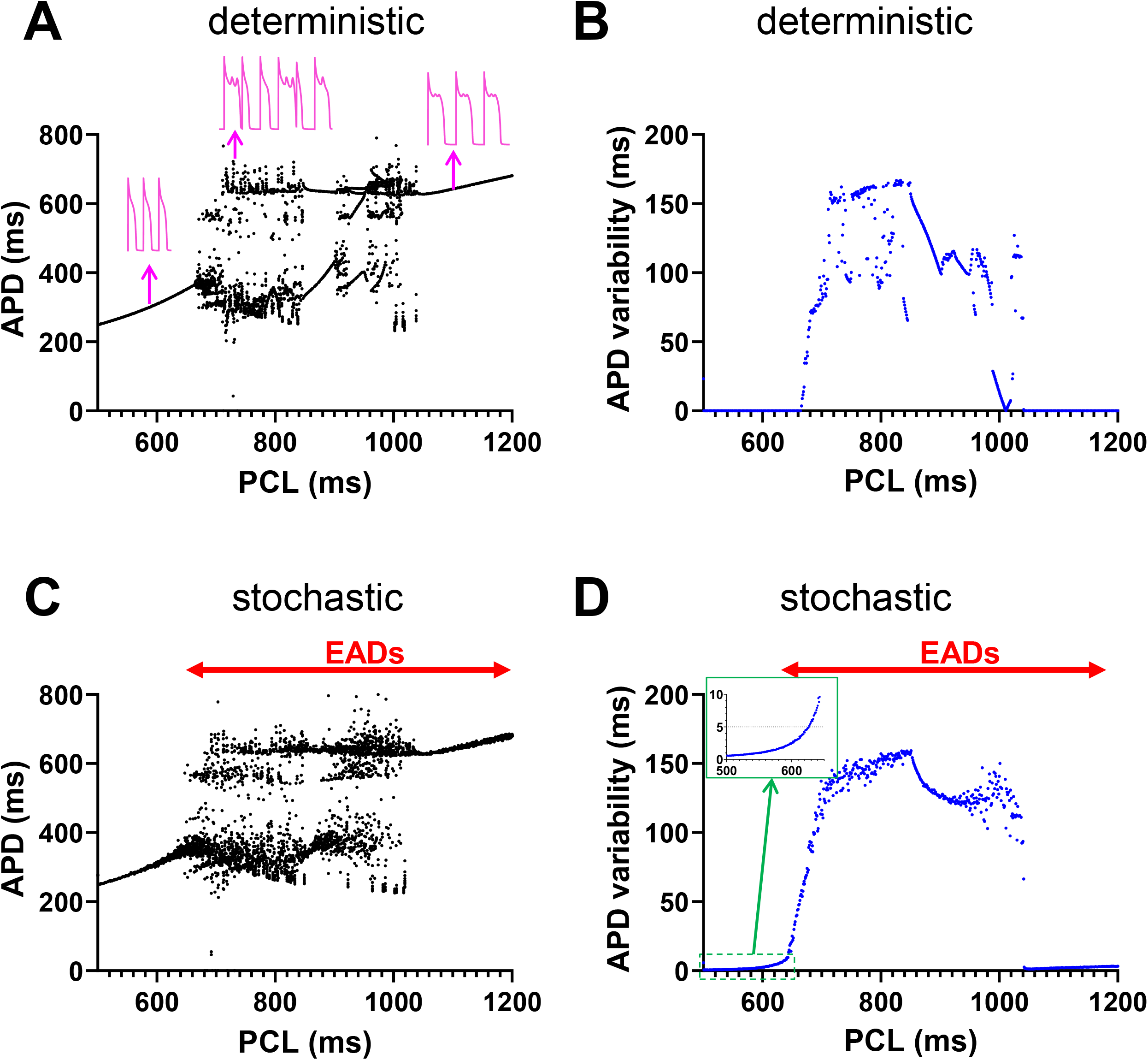
Stochastic noise affects APD variability. (A) APD vs PCL (deterministic model) (B) APD variability vs PCL (deterministic model). (C) APD vs PCL (stochastic model) (D) APD variability vs PCL (stochastic model). Inset: increase of APD variability in the pre-EAD regime. QTV > 5 ms (dashed line) often considered elevated arrhythmia risk.

Notably, we observed increased APD variability even without apparent EADs (Fig. 2D, inset). This observation is particularly important because it suggests that changes in APD variability, and consequently QTV on ECGs, may serve as an early indicator of arrhythmogenic EAD activity. Our study specifically investigates the regime just before the onset of EADs, where the AP is approaching dynamical instability but has not yet manifested apparent EADs. In this pre-EAD regime, we observe that stochastic fluctuations at the ion channel level are amplified by the underlying dynamical instability, resulting in increased APD variability. This amplification mechanism shown below provides a mechanistic link between observed APD variability and the propensity for EADs, even when EADs are not yet visible.

This increase in APD variability can be affected by various factors. One such factor is noise strength. Simulations with *N*=10,000 and *N*=1,000,000 show that larger noise (smaller *N*) leads to earlier increases in APD variability, well before EAD onset (Fig. 3). Nevertheless, in all cases, increased APD variability precedes the appearance of EADs.

**Figure 3.**
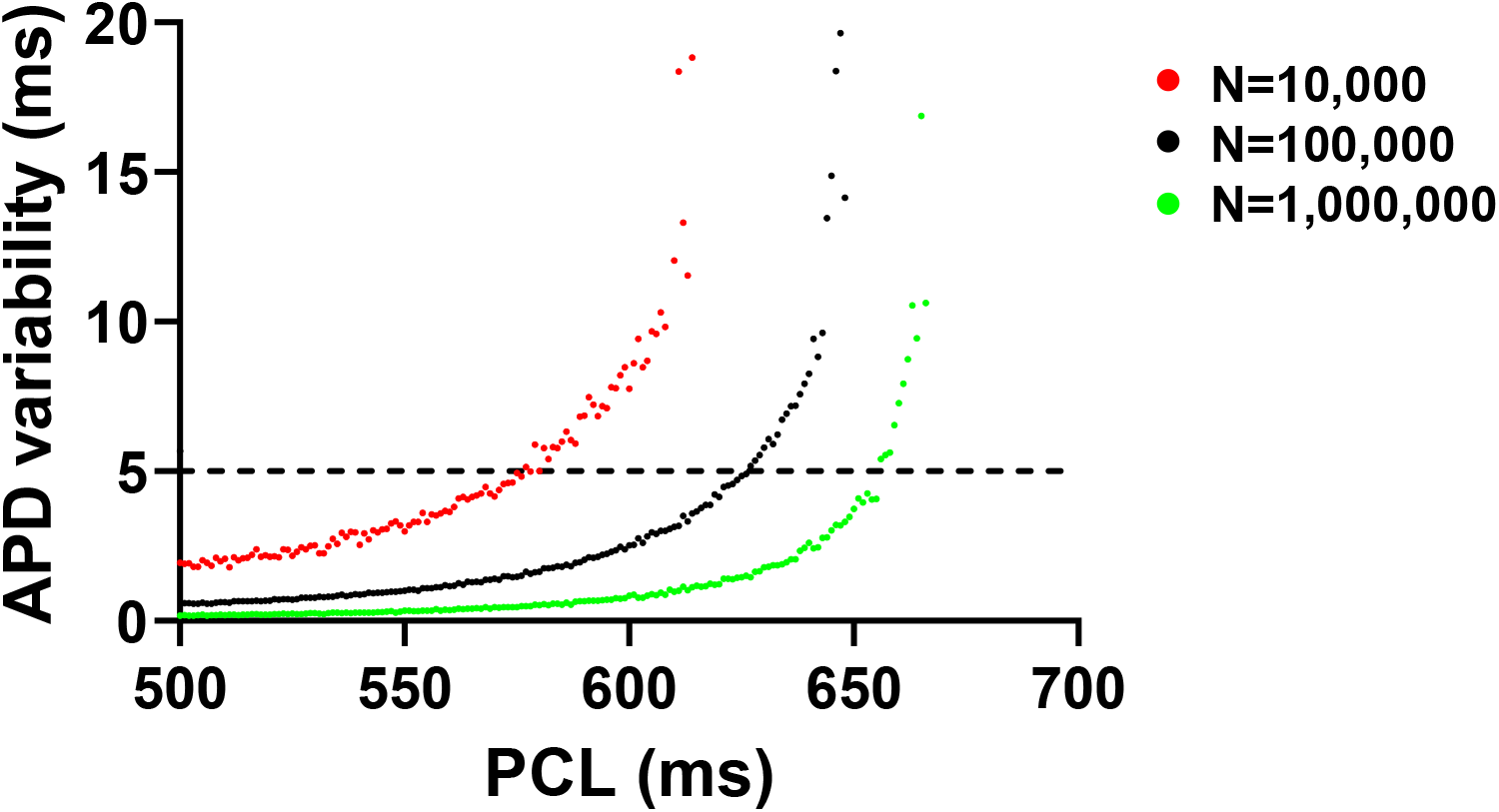
Effect of varying the noise strength on APD variability. APD variability as a function of PCL for different noise levels. Red circles: *N* = 10,000, Black circles: *N* = 100,000, Green circles: *N* = 1,000,000. As *N* decreases, the intrinsic noise strength increases, leading to APD variability emerging earlier and with a larger magnitude before EAD onset.

Our simulation results are quantitatively comparable to experimental data from rabbit myocytes (Fig. S2) (38). In experiments, the APD variability using 50 consecutive APs in healthy control rabbits ranged from 3∼7 ms, while in heart failure rabbits, the range widened to 3∼13 ms at PCL of 500 ms. In all cases, APs were normal (i.e., no alternans, EADs, or any other irregular APs).

In addition to noise strength, the sensitivity of APD to stochastic fluctuations depends on the model and its parameters, which affect the stability of the cell. This sensitivity becomes particularly pronounced under pathological conditions such as reduced repolarization reserve. In contrast, under healthy conditions with normal repolarization reserve, the AP is more stable and less sensitive to small perturbations and thus smaller variability (Fig. S3). In such cases, the pre-EAD amplification window is much broader than in the pathological condition. Moreover, disease progression or pharmacological interventions may shift or widen this window; individual variability can create different transition zones across patients; and normal daily heart rate fluctuations may intermittently cause QTV increases as patients enter or exit this vulnerable regime.

### APs become sensitive to small changes in the membrane voltage at longer PCLs

To understand the underlying mechanisms of increased APD variability without apparent EADs, we analyzed how the stability of APs changes with PCL (Fig. 4). To quantify the sensitivity of the system to stochastic noise, we systematically varied the stimulation current in the deterministic model. This approach allows us to probe the stability of the system in response to controlled perturbations, simulating the effect of small fluctuations that naturally occur at the channel level. By analyzing how small changes (i.e., initial membrane depolarization) affect APD, we can directly assess the degree of dynamical instability at different PCLs. We applied different stimulation currents ranging from 35 to 60 μA/μF (1 ms duration) to initiate APs. Smaller stimulation currents resulted in lower initial depolarization. Typically, this reduces the I_CaL_ and shortens the APD. As the stimulation current increases, I_CaL_ increases, and the APD becomes longer. However, excessive stimulation currents lead to excessively high initial membrane potentials, which reduce I_CaL_ due to a diminished driving force, consequently shortening the APD. The relationship between the initial depolarization level (resulting from varied stimulation currents) and APD is further illustrated in Fig. S4. These small changes in the initial membrane potential can lead to large APD changes depending on the stability. For example, chaotic EADs are highly sensitive to such changes (7-9), resulting in large variations in APD. Importantly, even with periodic APs, stability varies. At shorter PCLs, the cell exhibits greater stability, and variations in the stimulation current produce minimal changes in APD (Fig. 4A). In contrast, at longer PCLs, the cell becomes more sensitive to changes in membrane voltage, resulting in larger variations in APD for the same magnitude of stimulation variability, yet without EADs (Fig. 4B). This increased sensitivity at longer PCLs suggests that pacing frequency modulates the system’s stability and, consequently, APD variability (Fig. 4C). This subtle instability contributes to increased APD variability, which is potentially reflected in QTV on ECGs.

**Figure 4.**
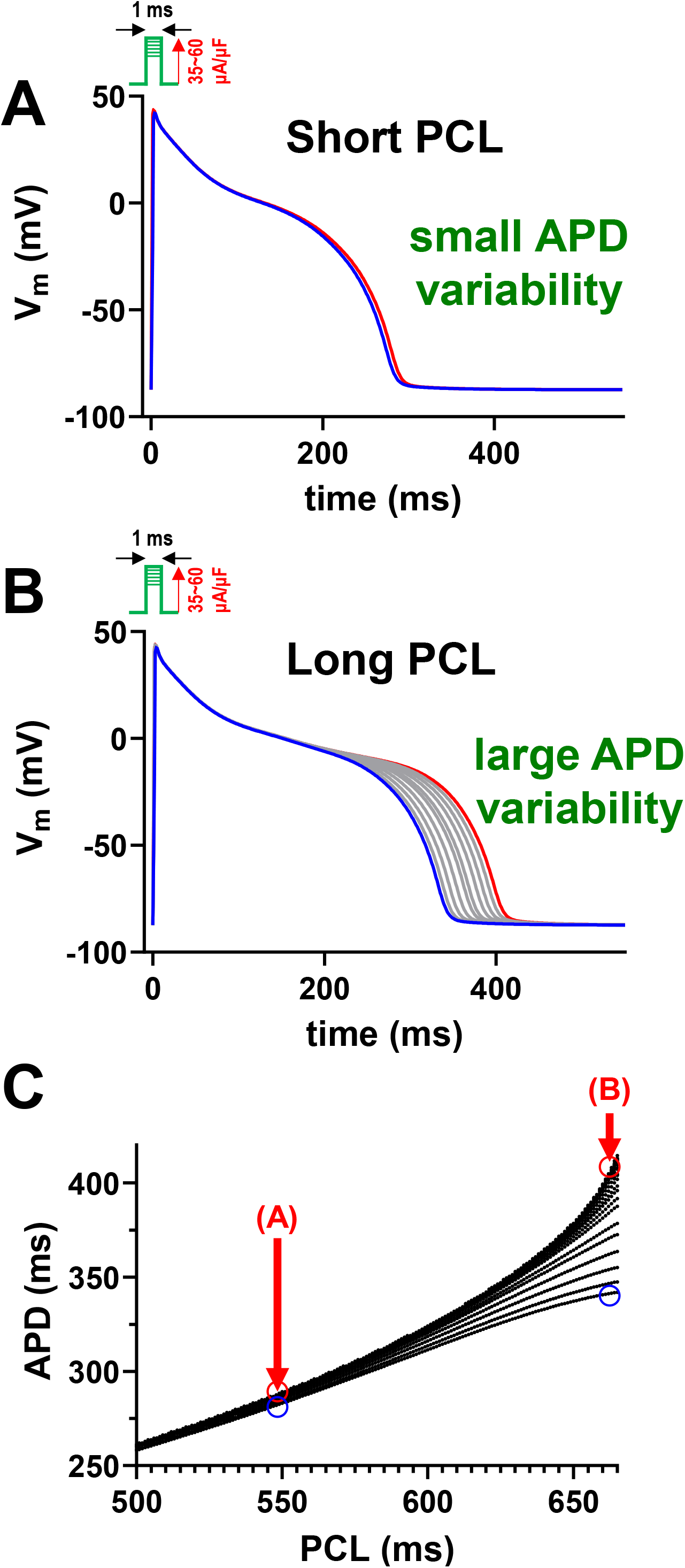
As the PCL approaches the onset of EADs, the membrane voltage becomes more sensitive to changes in the membrane voltage. (A) APs are less sensitive to changes in V_m_ at short PCL (PCL=550 ms) (B) APs are more sensitive to changes in V_m_ at long PCL (PCL=663 ms) (C) APD distribution and PCL

### EAD-generating basin of attraction strongly influences repolarization variability

To gain further insights, we analyzed the phase-plane of the *v-f* space, where *v* represents the membrane voltage and *f* denotes the voltage-dependent inactivation of the LTCC. In the detailed physiological model, the gating of the L-type Ca^2+^ channel is described by a 7-state Markov model. However, for simplicity and generalizability, we reduced it to a Hodgkin-Huxley type formulation, commonly used in many cardiac models such as the Luo-Rudy phase I (LR1) model (39). Following ref (9), the equations for the *v-f* space are

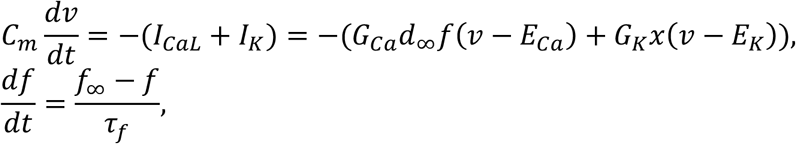

Where C_m_ is the membrane capacitance, G_Ca_ is the maximum conductance of LTCC, *d*_∞_ is the steady state of the activation of LTCC, *f*_∞_ is the steady state of the inactivation of LTCC, E_Ca_ is the reversal potential of I_CaL_, and G_K_ is the maximum conductance of the generic K current, E_K_ is the reversal potential of I_K_, and τ_*f*_ is the time constant of the *f* gate. The variable *x* represents the activation of I_K_, but is treated as a constant in this analysis. The details of the equations and parameters are shown in the appendix.

EADs occur when the cell state enters a specific region in the phase plane known as the basin of attraction, shown in green in Figs. 5A-C. This basin represents the set of initial conditions that lead to the reactivation of the L-type Ca^2+^ channel during the plateau phase of the APs. When the cell state is outside this basin, APs remain normal without EADs. Figs. 5A-C demonstrate that the initial proximity of the cell state to this basin critically influences APD variability. If the initial state is far from the basin, the variability remains low over time (Fig 5A). The spread of the membrane potential after 20 ms is within a 3mV range (−65 to –68 mV). Conversely, when the initial state is near the basin, increased variability over time was observed (Fig. 5C). The spread of the membrane potential after 20 ms is over a 63mV range (+2 to –61 mV). This phase-plane analysis highlights that APD variability is not simply random noise but is intimately linked to the underlying dynamical instability of the cardiac myocyte. The proximity to the EAD-generating basin of attraction dictates the cell’s susceptibility to perturbations and its resulting APD variability. This geometrical perspective, using phase-plane analysis, provides a powerful tool for visualizing and understanding the complex dynamics of EAD formation and APD variability. It suggests that subtle shifts in the cellular state, moving it closer to or further from the EAD basin of attraction, can have profound effects on electrical stability. Conversely, we can infer how close the system is to the EAD basin of attraction from the APD variability.

**Figure 5.**
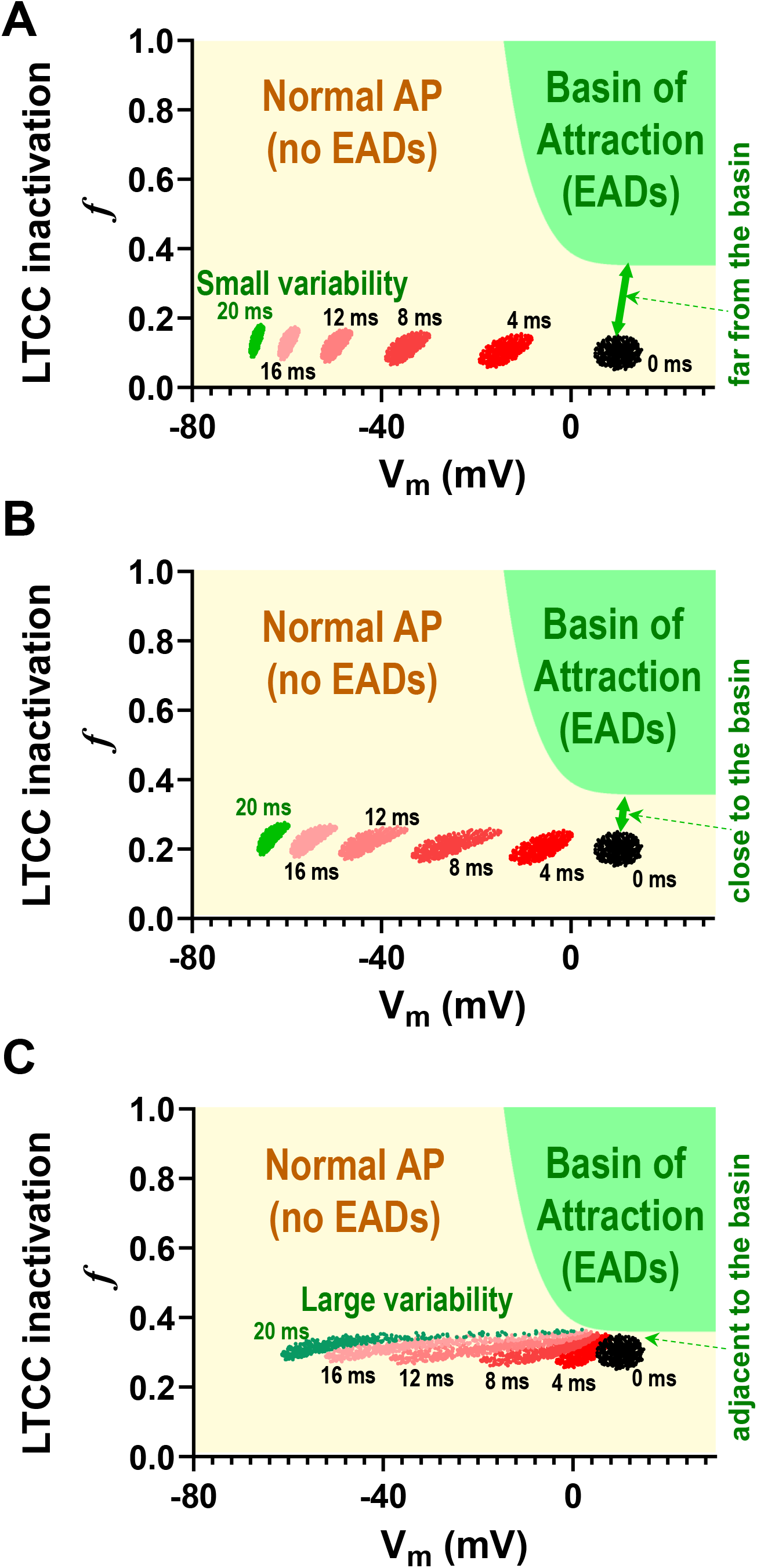
Variability increases as the cellular state moves closer to the basin of attraction. (A) Phase-plane trajectories when initial conditions are far from the EAD basin of attraction. Variability is limited in this case. (B) Variability increases as the cellular state moves closer to the basin of attraction. (C) Variability increases further as the cellular state moves even closer to the basin of attraction. In these phase-plane plots (v-f space), colors represent the temporal evolution of trajectories. Initial conditions (t=0 ms) are shown as a cluster of points (black), and their subsequent spread illustrates variability at later time points (red shades for 2-16 ms, green for 20 ms).

### Modulation of AP trajectories

We further analyzed trajectories in the *v-f-x* phase space, where *v* represents the membrane voltage, *f* denotes the inactivation of the LTCC, and *x* corresponds to the activation of a generic outward current (e.g., I_Ks_, I_Kr_) (9). In this analysis, *x* is a variable that follows the equation:

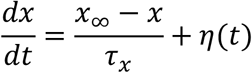

where *η*(*t*) is the noise term, which satisfies the following correlation function:

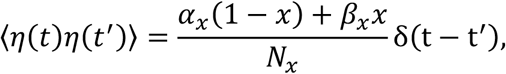

where *N*_*x*_ is the number of channels, α_*x*_ is the opening rate, and β_*x*_ is the closing rate. The resting state is indicated with “*” in Fig. 6A. The AP is initiated by a delta function that instantaneously shifts the voltage from the resting potential to +80 mV upon pacing.

**Figure 6.**
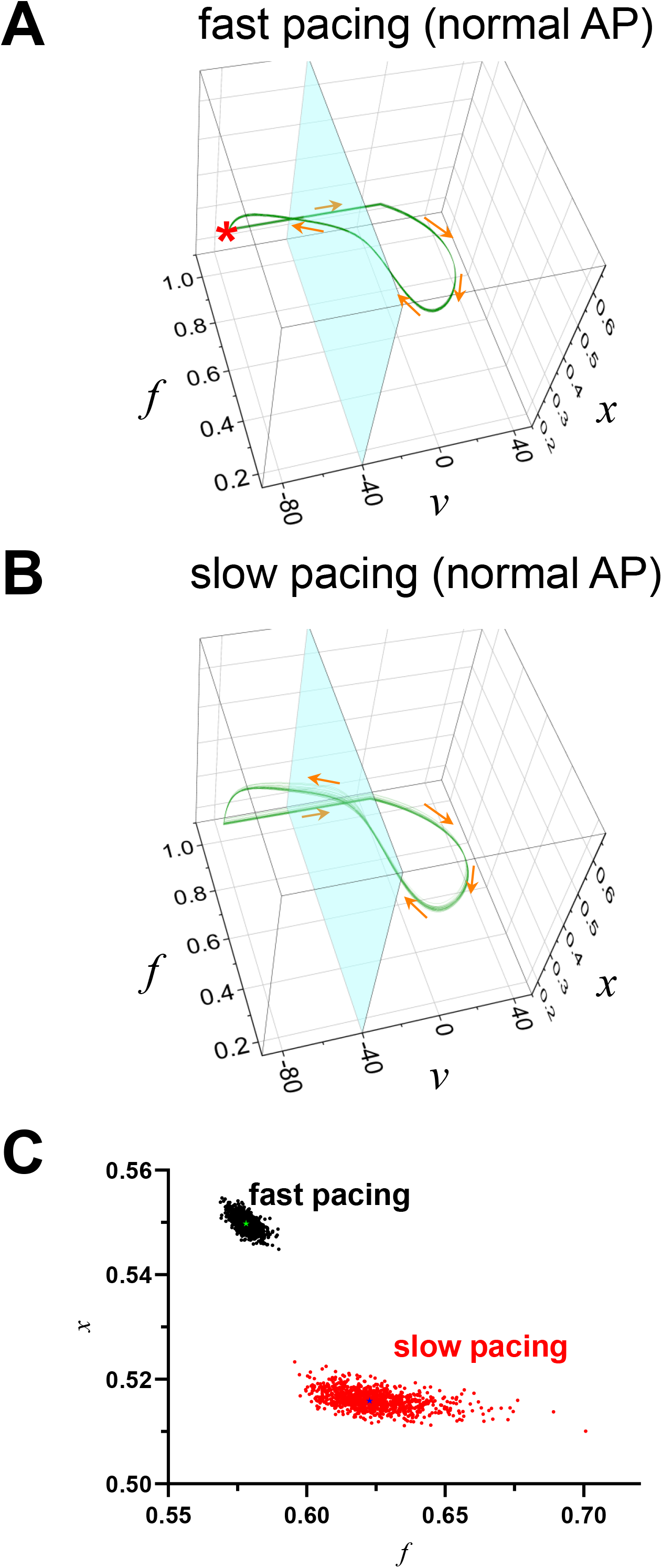
Dynamics in v-f-x space. *v* is the membrane voltage, *f* is the inactivation of LTCC, and *x* is the generic K current activation. In this model, the *x* gate is stochastic. (A) At fast pacing rates, the AP is stable. Therefore, stochastic noise has limited effects on the trajectory. (B) At slow pacing rates, the AP becomes less stable (still stable), and trajectories are highly affected by stochastic noise. (C) *f-x* plane at *v* = –40 mV. Black dots: fast pacing, Red dots: slow pacing. Green star indicates the deterministic trajectory for fast pacing, and Blue star indicates the deterministic trajectory for slow pacing.

When the pacing cycle length (PCL) is short (i.e., fast pacing), the AP is normal without EADs, and the trajectories remain closely clustered around the trajectory of the deterministic model (Figs. 6A and 6C black dots). However, as the PCL increases, the trajectories become more dispersed, leading to greater variability (Figs. 6B and 6C red dots). Fig. 6C shows the *f-x* plane at *v* = –40 mV (light blue plane in Figs. 6A and 6B). This plot demonstrates that variability increases at slower pacing (black dots to red dots), even in normal APs without EADs.

Fig. 7A shows that stochastic noise can alter the normal AP trajectory to generate an EAD: initially, the system follows the trajectory of a normal AP, but later, it transitions to a trajectory exhibiting an EAD (red loop). Conversely, Fig. 7B demonstrates the opposite scenario, where the trajectory initially follows the EAD path but later shifts to the normal AP trajectory (red line without loop). Fig. 7C presents two trajectories –one corresponding to a normal AP and the other to an EAD. In this case, the system alternates between these two trajectories. However, in some instances, stochastic noise causes the system to transition from one trajectory to the other (red line), resulting in a phase flip of the alternating EADs.

**Figure 7.**
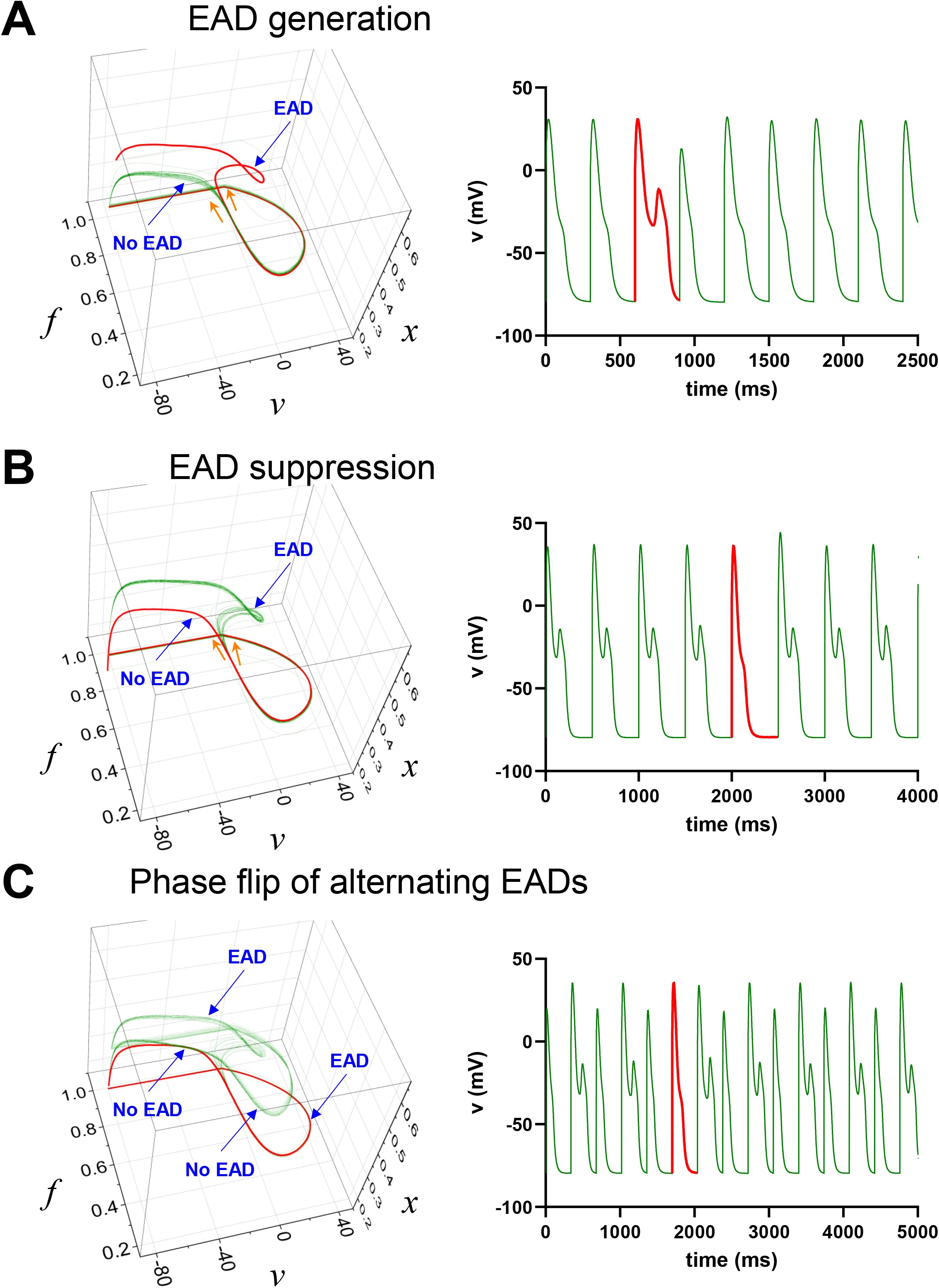
Stochastic noise and EAD generation and suppression. (A) If the PCL is close to the EAD regime, normal AP can become EAD due to stochastic noise. The red line shows the trajectory that entered the EAD trajectory. (B) EADs can be suppressed due to stochastic noise. (C) Stochastic noise causes a phase flip of the alternating EADs

### Increased QTV in 2D tissue precedes EADs

To assess whether the observed single-cell APD variability translates to the tissue level, where electrotonic coupling can reduce variability, we performed 2D tissue simulations (Fig. 8). Pseudo-ECGs were recorded and QTV was calculated (Figs. 8A and B). Consistent with our single-cell findings, QTV in the 2D tissue model increased as the PCL approached the onset of EADs (Fig. 8C). Near the EAD threshold, fluctuations were progressively amplified and this amplification persisted until EADs appeared, at which point QTV dramatically increased, exceeding 100 ms even in the absence of stochastic noise (Fig. 2B). The magnitude of this pre-EAD QTV was greater than the clinically relevant range typically associated with increased arrhythmia risk (> 5 ms) (40). These results support our central hypothesis that increased QTV precedes the onset of EADs and is observable even in electrotonically coupled tissue.

**Figure 8.**
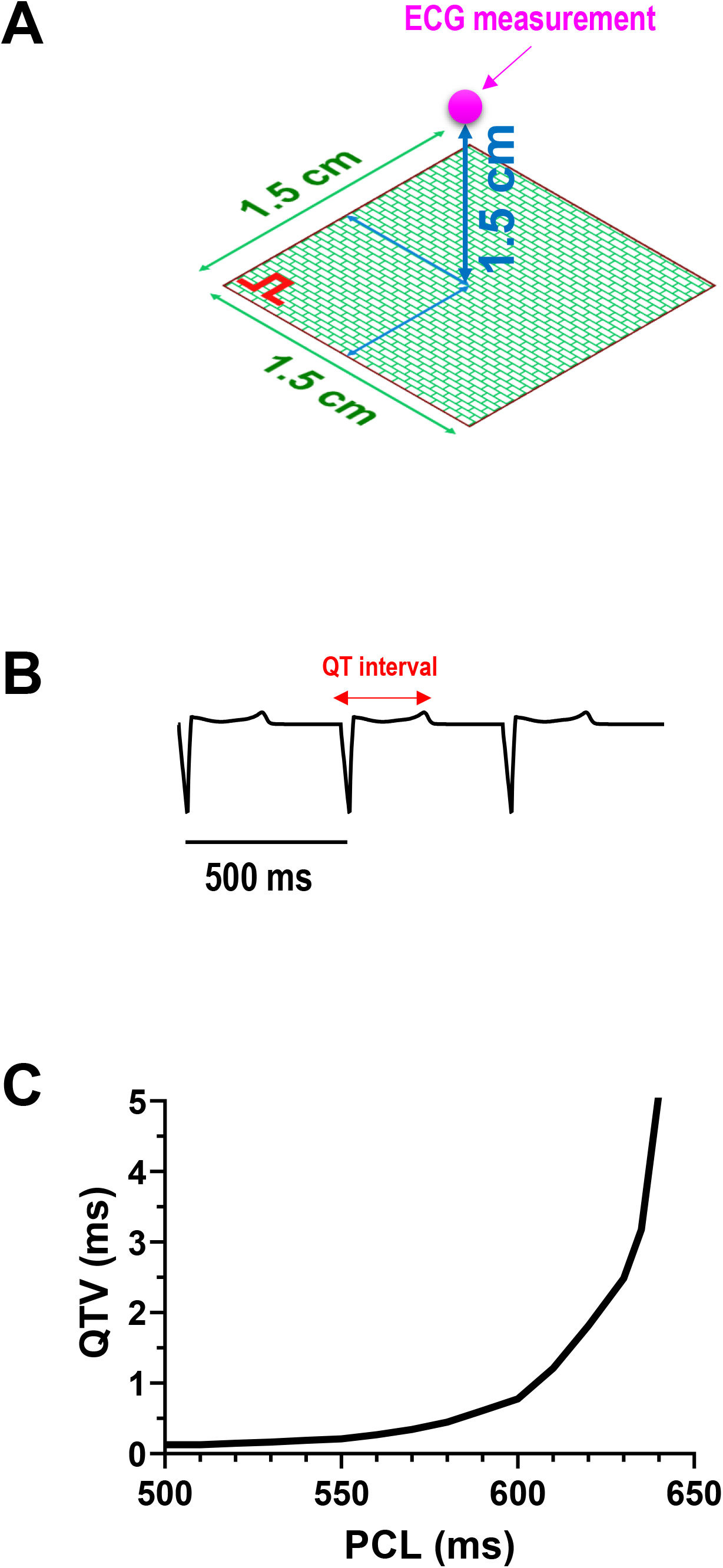
Increased QTV precedes EADs in 2D tissue simulations. (A) The tissue was modeled as a 100 × 100 grid (1.5 cm × 1.5 cm) of coupled cells. Pacing was applied at one corner of the tissue. Pseudo-ECGs were recorded at the location indicated by the magenta circle. (B) Representative pseudo-ECG trace at PCL=500 ms. (C) QTV as a function of PCL.

### Underlying mechanisms of QTV and their clinical relevance

In a recent study, we demonstrated that APD variability increases before the onset of alternans due to increased instability, which occurs as the PCL becomes shorter (faster pacing rates) (29). In contrast, EADs resulting from membrane potential instability are typically observed at longer PCLs (slower pacing rates). In this study, we showed that APD variability increases as PCL becomes longer, which is driven by the membrane potential becoming unstable (Fig. 4). In both scenarios, the instability arises from the inherent nonlinearity of the cardiac AP dynamics, underscoring its critical role in modulating APD variability and thus QTV. This suggests that increased QTV can be a general marker of cardiac electrical instability across different pacing regimes, albeit reflecting distinct underlying mechanisms and potentially different pro-arrhythmic risks. In fact, both cases have been experimentally observed (40,41).

These findings have important implications for the understanding of QTV and its clinical significance. While QTV has been associated with an elevated risk of arrhythmias (15-23), its underlying mechanisms are not fully understood. Our results suggest that increased APD variability, even in the absence of apparent EADs, could reflect subtle dynamical changes in the heart that precede arrhythmogenesis. This “pre-arrhythmic state,” characterized by increased QTV without overt arrhythmias, may represent a critical window for early intervention and risk stratification. For instance, in patients with risk factors for sudden cardiac death, such as heart failure or ischemic heart disease, monitoring QTV might provide an early warning sign of increased electrical instability, prompting more aggressive preventative strategies or closer monitoring. Similarly, in the context of drug-induced QT prolongation, an increase in QTV might signal heightened EAD risk and the need to adjust medication regimens or consider alternative therapies. Furthermore, even among healthy individuals, variations in QTV may serve as an early indicator of arrhythmia risk.

### Limitations and Future Directions

In this study, we focused on EADs due to the reactivation of LTCC. EADs can also be caused by spontaneous Ca^2+^ releases from the sarcoplasmic reticulum (42-44); however, these phenomena were not incorporated into our model. Capturing these events poses significant mathematical and computational challenges due to their spatial complexity. Future studies should aim to integrate detailed subcellular Ca^2+^ handling into the model to explore the interplay between intracellular Ca^2+^ dynamics and EAD-related QTV. In addition, our investigation was conducted using a single-cell computational model of a rabbit ventricular myocyte. Rabbit ventricular electrophysiology and Ca^2+^ handling are similar to human (45), and while some details differ, our conclusions are likely to be broadly valid in human. This model is physiologically detailed at the cellular level allowing controlled exploration of cellular mechanisms, but cannot fully capture the complexities of the intact heart. In fact, we measured APD variability at the single cell level, but this is not equivalent to QTV observed on the surface ECG. While our 2D tissue simulations (Fig. 8) demonstrate that the core phenomenon of increased QTV near the EAD threshold persists in coupled tissue, these simulations are still a simplification. In the intact heart, QTV is influenced by additional factors including transmural and spatial heterogeneities in electrophysiology, cell-to-cell conduction, and the propagation patterns of activation and repolarization throughout the ventricles. Future studies should incorporate more detailed multicellular simulations and anatomically realistic tissue-level models to investigate how QTV and EADs manifest at the tissue and whole-heart levels.

Lastly, the direct clinical translation of our findings requires further validation in experimental and clinical settings. Future research should focus on correlating QTV measures in vivo with cellular electrophysiological properties and EAD susceptibility in animal models and human subjects. Investigating the pharmacological modulation of QTV and EAD susceptibility based on our mechanistic insights could also lead to novel therapeutic strategies for arrhythmia prevention. Indeed, previous studies have shown that various agents can suppress or increase both QTV and arrhythmogenic events, such as EADs and TdP, simultaneously (46-50). Our study demonstrated that both increased QTV and EADs are driven by dynamical instability. Therefore, it would be valuable for future studies to investigate whether the suppression of QTV and arrhythmogenic events by such agents is a result of the stabilization of this dynamical instability.

## Conclusions

This study demonstrates that increased APD variability, and thus increased QTV, can precede the manifestation of apparent EADs. As the cellular state approaches the onset of EADs, APD variability increases significantly (Fig. 2). While alternans-associated APD variability increases with faster pacing (29), EAD-associated APD variability increases with slower pacing. This distinction enables differentiation between alternans-related and EAD-related arrhythmic risk. Our findings suggest that increased QTV reflects subtle dynamical changes in the heart (Figs. 5 & 6), serving as a predictive indicator of arrhythmic risk. This underscores the importance of both stochastic and nonlinear factors in assessing cardiac instability and provides insights for improving arrhythmia risk stratification and prevention.

## ABBREVIATIONS

AP: action potential
APD: action potential duration
Ca^2+^: calcium
Na^+^: sodium
QTV: QT interval variability
ECG: electrocardiogram
HRV: heart rate variability
PCL: pacing cycle length
EAD: early afterdepolarization

## Author Contributions

D.S. performed mathematical modeling and computer simulations. D.S. processed and analyzed the simulation data. D.S., B.H., C.M.R and D.M.B. interpreted the data and wrote the manuscript. All authors approved the final version of the manuscript.

## Acknowledgments

Supported by NIH grants (P01-HL141084, R01HL142282 DB; R01HL149349 DS and DB; R01HL111600 CMR) and American Heart Association grant 23CDA1051603 BH.

---

**Figure S1.**
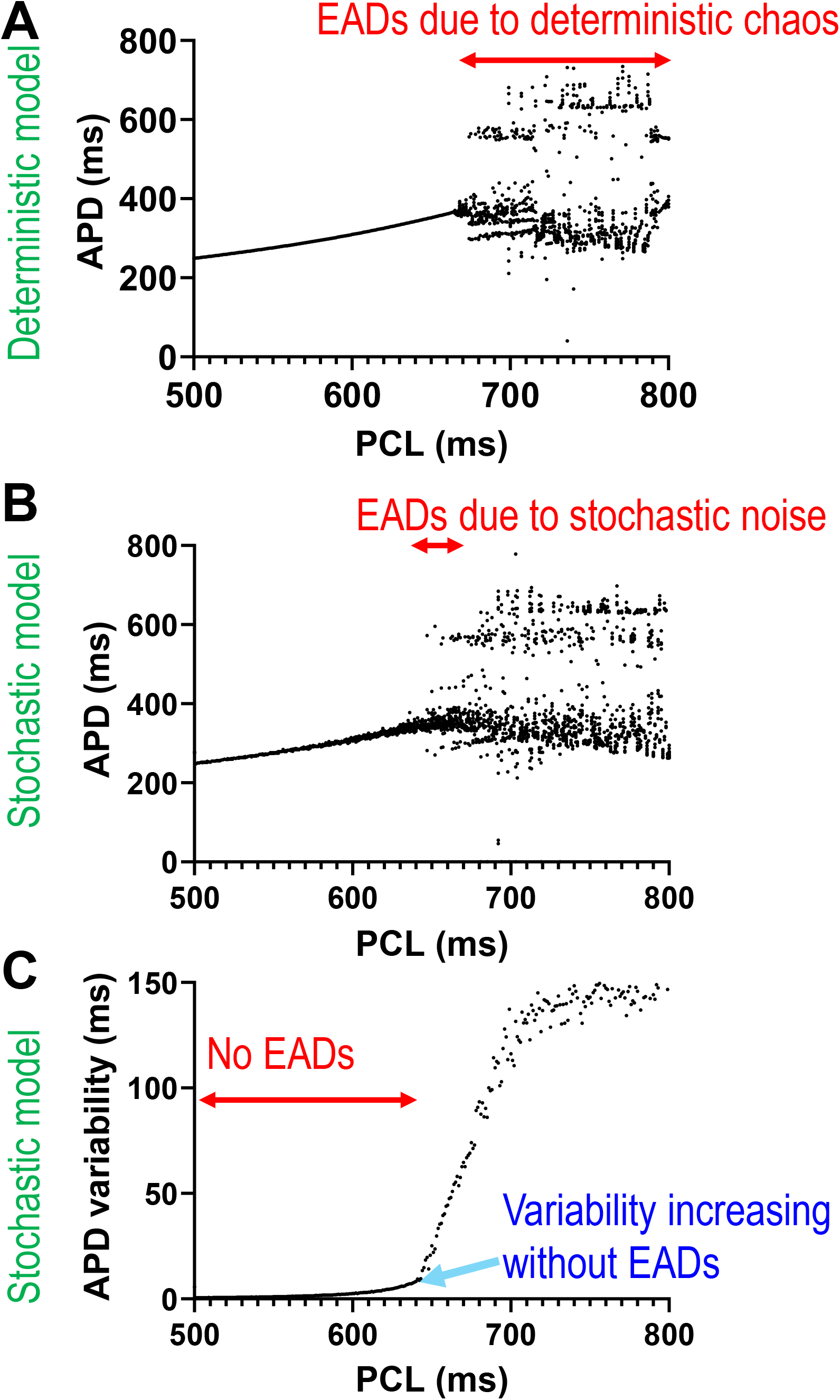
Detailed view of the data presented in Figure 2, focusing specifically on the PCL range of 500 ms to 800 ms. (A) APD (from Fig. 2A) for PCLs between 500 and 800 ms. Large variability is due to deterministic chaos. (B) APD (from Fig. 2C) for the same PCL range. Large variability is due to stochastic EADs. (C) APD variability (from Fig. 2D) for the same PCL range. Variability increases without EADs.

**Figure S2.**
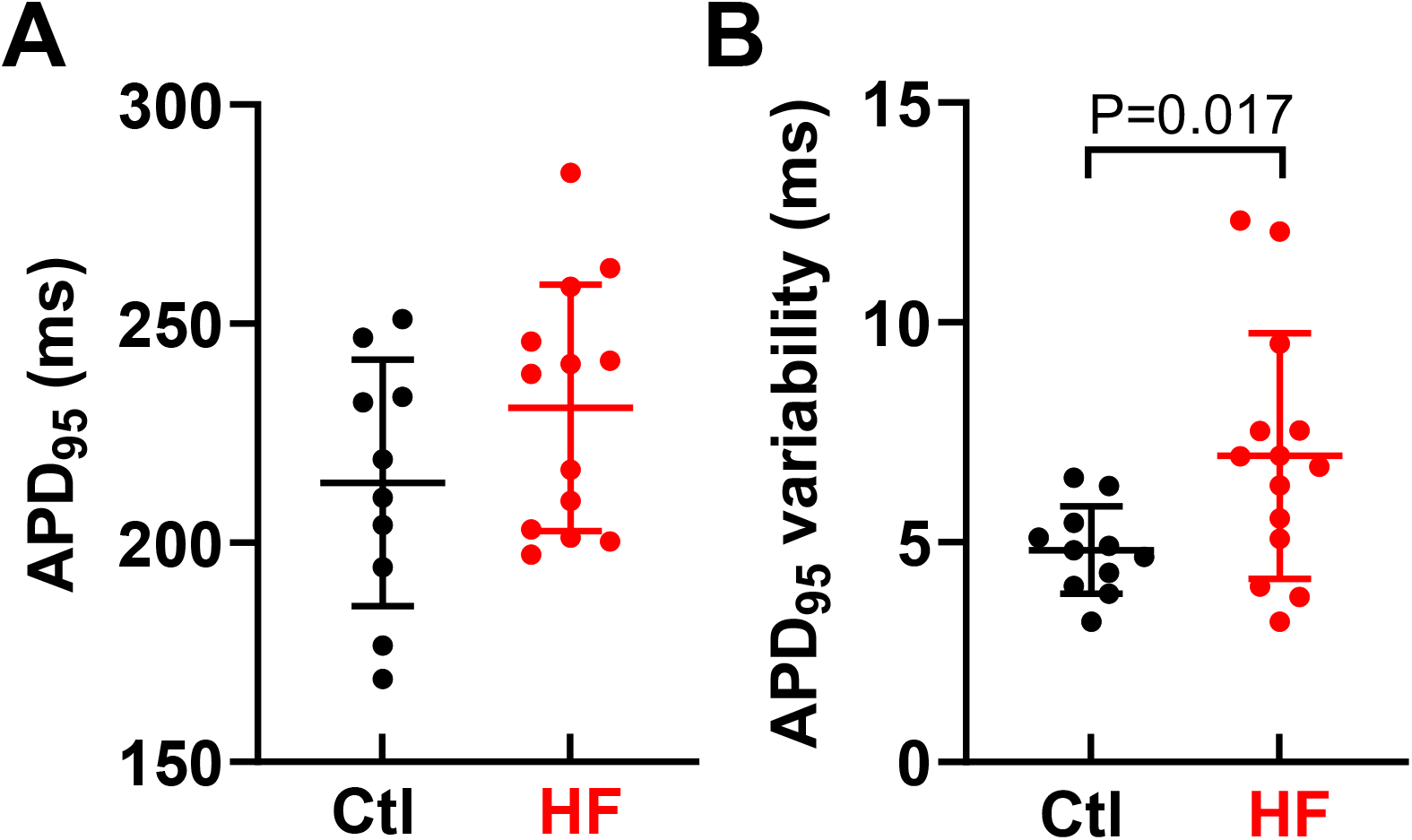
APD variability in rabbit ventricular myocytes. Data are from Ref. (38). In that study, variability was calculated as the short-term variability of APD, whereas in the present study, we quantify APD variability using the standard deviation of APD. Although we used the same dataset, the recalculated APD variability values are slightly different from the values reported in Ref. (38). (A) In healthy control rabbits, APD variability ranged from 3 to 7 ms. (B) In heart failure rabbits, it ranged from 3 to 13 ms.

**Figure S3.**
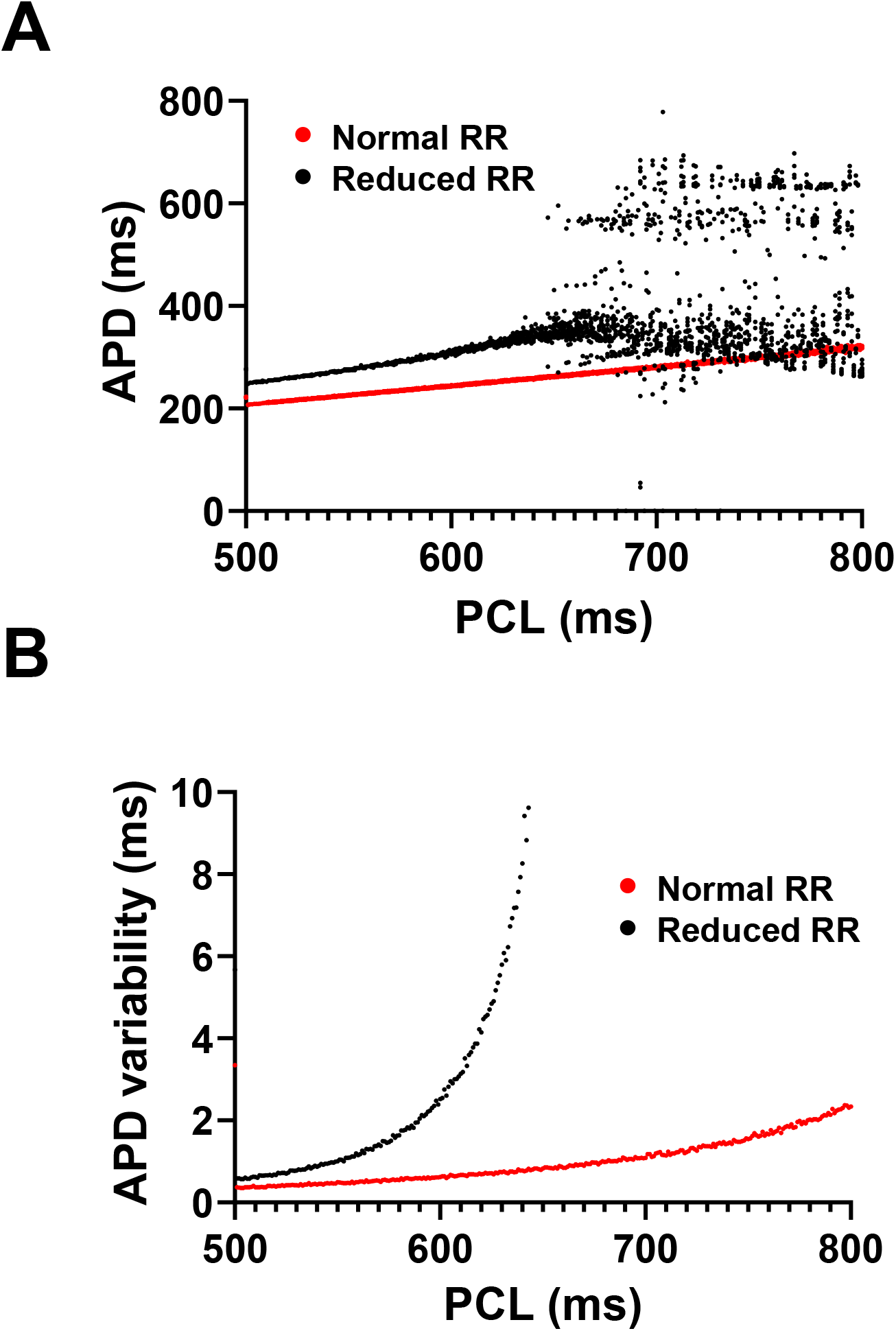
APD variability for normal repolarization reserve AP model. Black: reduced repolarization reserve (same as Fig 2), Red: normal repolarization reserve. (A) APD vs. PCL. (B) APD variability vs. PCL.

**Figure S4.**
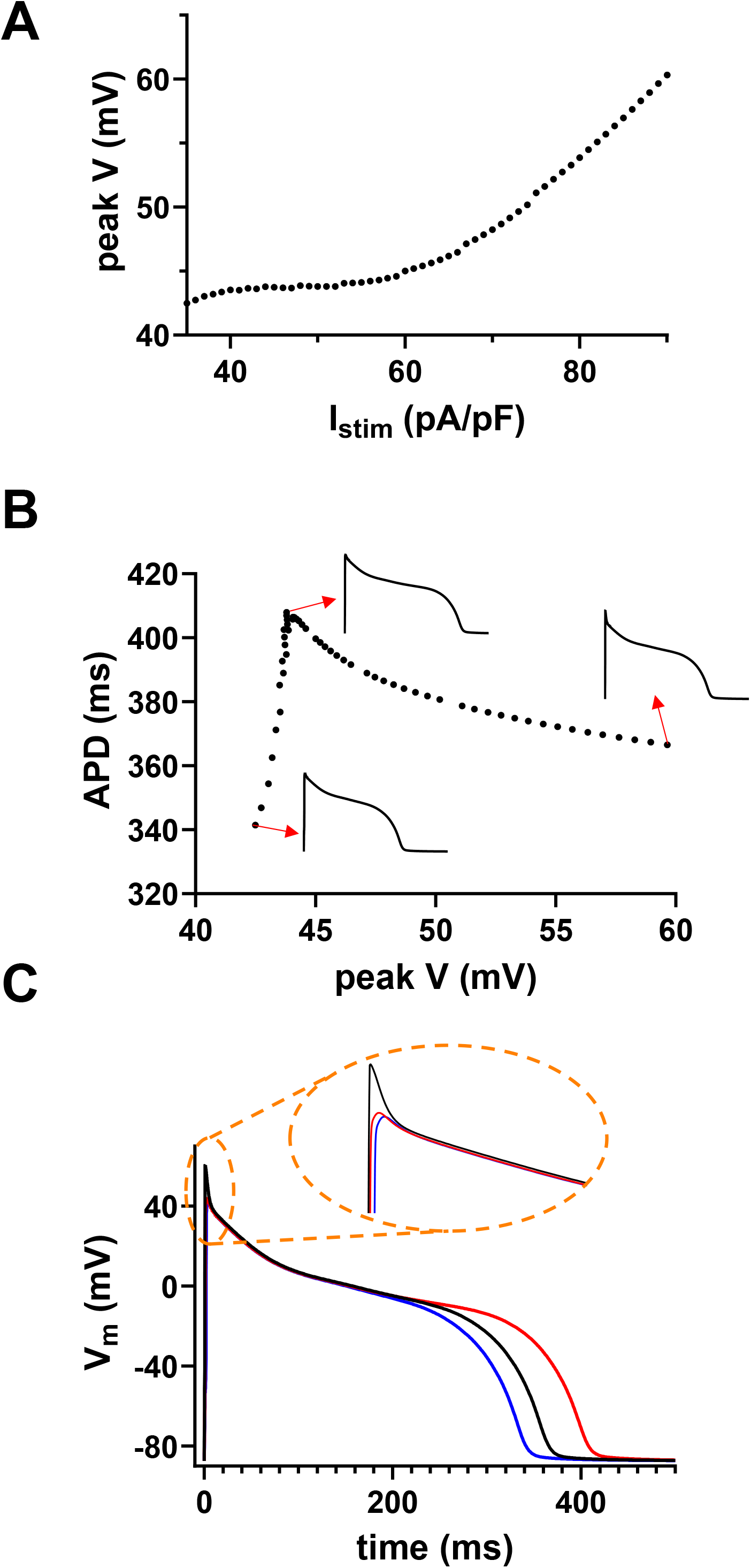
Relationship between initial depolarization level and APD. (A) Variations in stimulation current strength lead to different initial membrane depolarization levels (B) Changes in the initial membrane depolarization level result in a biphasic APD response. (C) AP traces. Blue: I_stim_= 35 μA/μF, Black: I_stim=_ 90 μA/μF, Red: I_stim_= 52 μA/μF.

